# Decision making in auditory externalization perception: model predictions for static conditions

**DOI:** 10.1101/2020.04.30.068817

**Authors:** Robert Baumgartner, Piotr Majdak

## Abstract

Under natural conditions, listeners perceptually attribute sounds to external objects in their environment. This core function of perceptual inference is often distorted when sounds are produced via hearing devices such as headphones or hearing aids, resulting in sources being perceived unrealistically close or even inside the head. Psychoacoustic studies suggest a mixed role of various monaural and interaural cues contributing to the externalization process. We developed a model framework for perceptual externalization able to probe the contribution of cue-specific expectation errors and to contrast dynamic versus static strategies for combining those errors within static listening environments. Effects of reverberation and visual information were not considered. The model was applied to various acoustic distortions as tested under various spatially static conditions in five previous experiments. Most accurate predictions were obtained for the combination of monaural and interaural spectral cues with a fixed relative weighting (approximately 60% of monaural and 40% of interaural). That model version was able to reproduce the externalization rating of the five experiments with an average error of 12% (relative to the full rating scale). Further, our results suggest that auditory externalization in spatially static listening situations underlie a fixed weighting of monaural and interaural spectral cues, rather than a dynamic selection of those auditory cues.

## Introduction

For a successful interaction with the environment, influential theories suggest that the brain’s primary objective is to infer the causes of its sensory input by creating an internal model of the environment generating expectations about the incoming cues [1]. These theories convince by examples coming from various areas of sensory perception and higher-order cognitive functions [2]. The underlying perceptual decision making is usually based on multiple, simultaneously accessible cues. Various decision strategies such as weighted sums or cue selection have been put forward in the context of visual search tasks [3] but remained an unresolved topic in spatial hearing [4]. The goal of this study is to test those concepts on a particularly puzzling perceptual phenomenon of spatial hearing, namely, the collapse of perceptual externalization (or distal attribution) [5], which denotes the inability to associate sensations with external objects [6]. Auditory externalization constitutes a critical aspect of spatial hearing because it can be easily disrupted when listening to sounds, e.g., via headphones or other hearing devices, that do not accurately represent the spatial properties of the listener’s natural acoustic exposure [7–9].

The spatial properties of the sound arriving at the two ears are manifold and so are the cues that can be used for spatial inference [10]. Many studies aimed to identify in particular spectral cues affecting the perceptual externalization of sound sources [7]. These cues can be categorized as either monaural or interaural. Monaural cues include the overall sound intensity, known as a dominant relative distance cue [11], or the spectral shape, known to be crucial for sound localization beyond the horizontal plane [12–14]. Interaural cues include the interaural intensity difference (IID), interaural time difference (ITD) and/or interaural coherence (IC), which are well known to be crucial for sound localization within the left-right dimension [15, 16] but may also affect spatial hearing in other dimensions [17, 18]. Moreover, cues from both categories may potentially be evaluated on a frequency-specific or broadband manner. Besides the multitude of potential cues, a general problem is also that many of those cues co-vary and that it has never been investigated which perceptual decision strategy may underlie the listeners’ response behavior in those psychoacoustic tasks.

While the cue-based encoding can be considered as trivial, probably already happening before reaching the auditory cortex [19], more complex structures are required to combine the cues to form a decision stage yielding the final percept of externalization. Cortical representations contain an integrated code of sound source location [20–22], but also retain independent information about spatial cues [23–26], allowing the system to decide how likely they would have arisen from one or multiple sources [26], or would have arisen from an external source, i.e., manifesting as a final percept of auditory externalization. The perceptual decision strategy used by the auditory system to form externalization is unclear yet. Here, we tested two general types of decision strategies.

The first potential strategy follows the idea that based on exposure statistics our brain has established a rather fixed weighting of information from certain spatial auditory cues in order to perceive an externalized auditory event – basically independent of the listening context. This can be represented as a static weighted-sum model (WSM, see Fig. 1d) that scales the expectation errors obtained for a single cue with weights adjusted based on long-term listening experience. Once the weights are settled, no further adaptation is considered. Such a static WSM is often used to merge neural processing of multiple cues, with examples ranging from a lower, peripheral level such as neural binaural processing [27] over higher cortical levels integrating visual orientation cues [28] to even more abstract levels such as the representation of person identities [29].

**Fig. 1.**
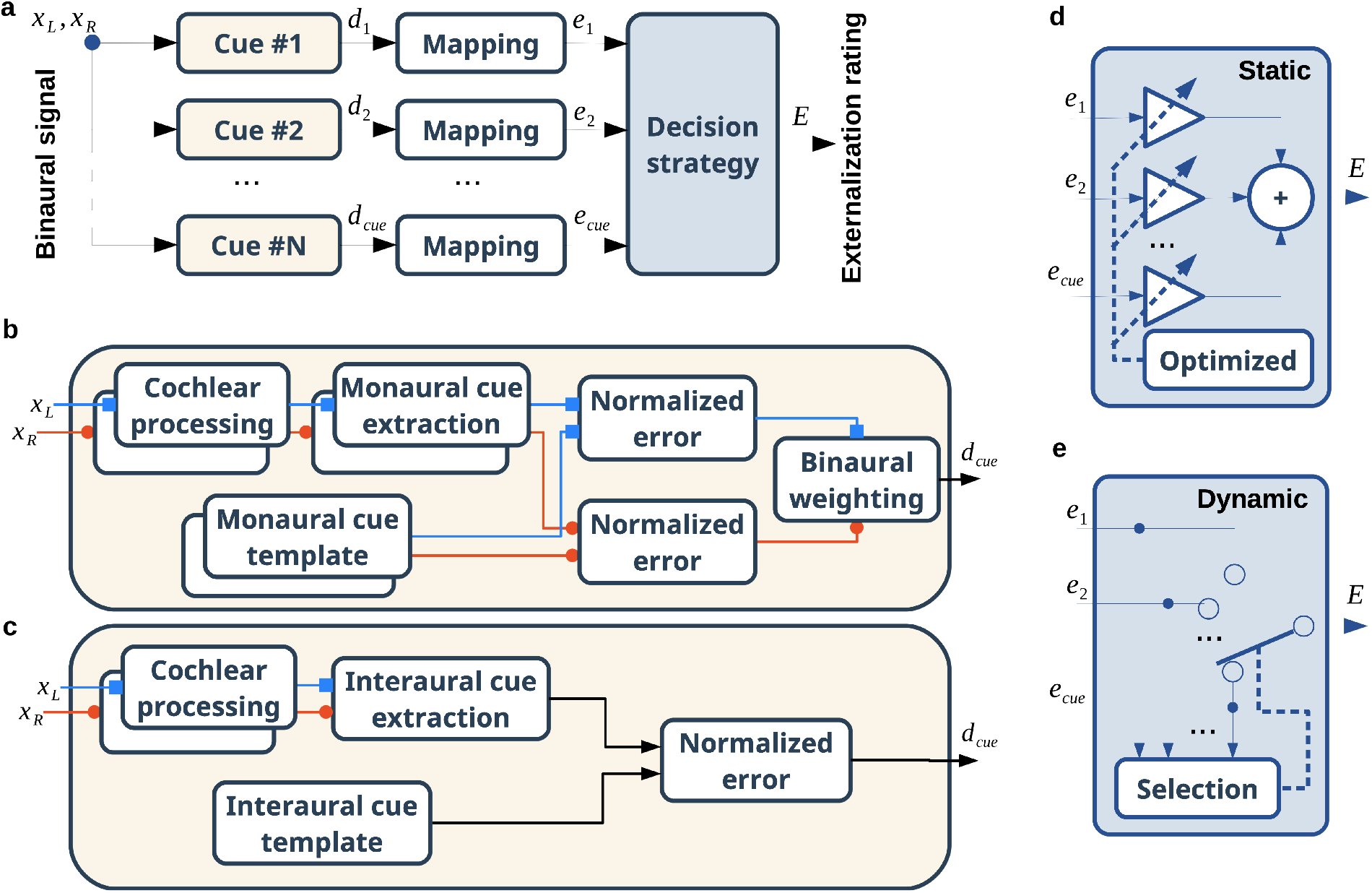
General structure of the sound externalization framework. **a**, Processing of the binaural signal x_L_, x_R_ resulting in a cue-based error d_cue_, a stage mapping the cue-based error to a normalized externalization rating e_cue_, and a decision stage combining the cue-specific ratings to a final externalization rating E. **b**, Processing of the binaural signal to calculate a single-cue error based on monaural cue templates. **c**, Processing based on interaural cue templates. **d**, Decision stage based on a static weighted-sum model (WSM). **e**, Dynamic decision stage based on the selection of a cue-specific rating. Minimalist (LTA), conformist (MTA), and perfectionist (WTA) approaches were considered for selection (see text for further explanation).

The second potential decision strategy is of selective nature and has been promoted, for instance, in the context of visual search [30, 31] or audio-visual dominance [32]. The idea is that, depending on the incoming stimulus, our brain selects the one of the externalization-related cues that fulfills or breaks one or most of the listener’s prior expectations (see Fig. 1e). We considered three variants of such a selection strategy. First, a minimalist approach would assume that a sound can be externalized if at least one cue promotes externalized perception by matching the listener’s expectations. We implemented this as a winner-takes-all (WTA) strategy that considers the largest externalization rating (minimum expectation error across all individual cues) as the total externalization rating. Second, the contrary is the perfectionist approach, which requires all of the cues to promote an externalized percept – the cue breaking its expectation the most will dominate the perception. This is implemented as the loser-takes-all (LTA) strategy, which considers the smallest externalization rating (maximum expectation error across cues) as the total rating. Third, the conformist approach is intermediate to these two extremes and selects the cue-specific expectation error being most consistent to all others. We implemented this as a median-takes-all (MTA) strategy, which considers the median across all single-cue-based ratings as the total externalization rating.

Here, we propose a model framework allowing us to disentangle those decision strategies potentially involved in the multi-cue-based task of auditory externalization. We used that framework to simulate five experiments from four representative psychoacoustic externalization studies [17, 33–35], all focusing on the effects of spectral distortions under spatially static listening conditions, but differing considerably in manipulation method and bandwidth. The model represents each listener’s expectations for perceptual inference as internal templates and uses them for comparison with incoming cue-specific information. The decision strategies were not only compared for the whole set of cues, but also for a successively reduced set, in order to address the potential redundancy between them. Published work that was conducted in parallel to the here presented modeling study built upon preliminary results from the present work and extended it by also incorporating reverberation-related cues [36].

## Materials and Methods

### Structure of the model mapping signals to externalization ratings

The model flexibly comprises a variety of cues as summarized in Tab. 1. We considered various cues that have been associated with externalization or distance perception of anechoic sounds: monaural spectral shape (MSS) [12, 37], interaural spectral shape (ISS) [17], the difference in broadband monaural intensity (MI), the monaural spectral standard deviation (MSSD) [34, 38], the spectral standard deviation of IIDs (ISSD) [39], the broadband interaural coherence (IC) [40], and the inconsistency between ITD and IID (ITIT). While most of these cues were directly considered as factors in previous experiments being modeled here, the ITIT has up to our knowledge never been quantified in the context of sound externalization. Our rationale behind it was that if the distinct combination of ITD and IID were used for the localization of nearby sources [18], then counteracting deviations for those two cues should distort the process of auditory externalization. The deviations in broadband ITD or broadband IID were not evaluated in isolation because it has been previously reported that offsets applied to either of those cues hardly affected externalization perception [16, 41].

**Tab. 1.**
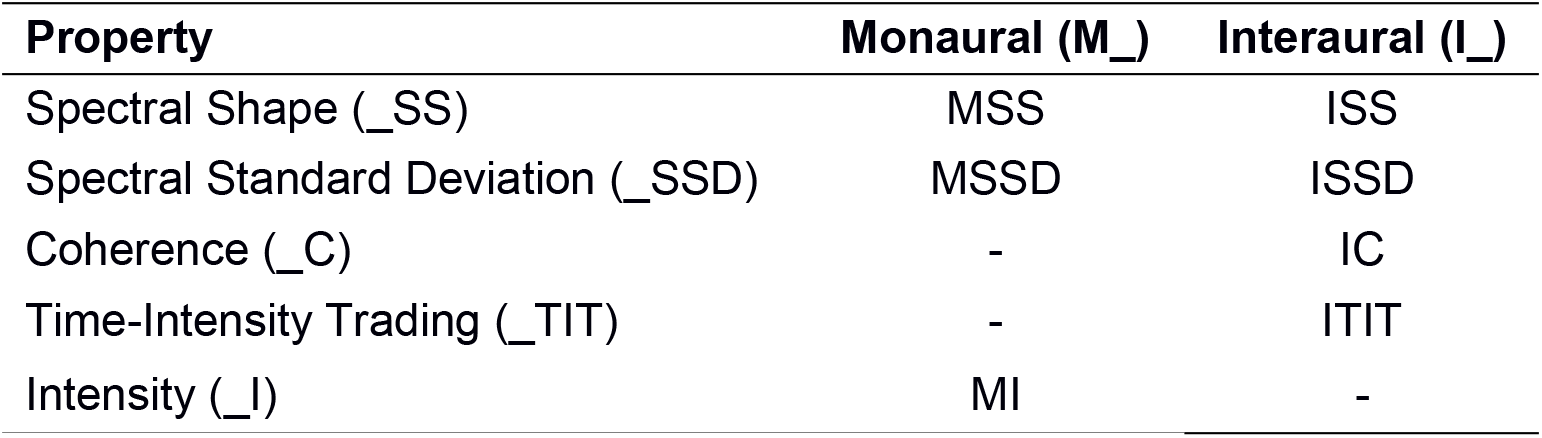
Potential auditory externalization cues considered for evaluation.

Depending on the considered cue, slightly different model architectures were required for processing either monaural (Fig. 1b) or interaural cues (Fig. 1c). Both architectures follow a standard template matching procedure [7, 42–44] as they take a binaural signal as an input and simulate externalization ratings as an output after performing comparisons with internal cue templates. Those templates were derived from listener-specific head-related transfer functions (HRTFs) or binaural room impulse responses (BRIRs), depending on the modeled experiment.

The model framework (Fig. 1a) consists of 1) cue-processing stages, each of them encoding the incoming binaural signal, [*x*_*L*_, *x*_*R*_], into a single-cue expectation error, *d*_*cue*_, 2) stages mapping that single-cue error to a normalized externalization rating, *e*_*cue*_, and 3) a decision stage combining the single-cue based ratings into the final externalization rating, *E*. The mapping from *d*_*cue*_ to *e*_*cue*_ was scaled individually for each experiment, unless otherwise stated.

### Error metrics

For the MI cue, the overall level differences were considered as:

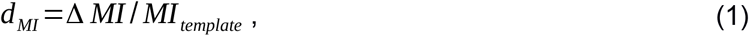

with Δ *MI* =|*MI*_*target*_ − *MI*_*template*_| denoting the difference in root mean square (RMS) levels (in dB) of the incoming broadband signals. Differences smaller than ±1 dB were considered to be below the just-noticeable difference [45] and thus set to zero. The error based on MI, *d* _*MI*_ was calculated for each ear separately and then averaged across both ears.

The IC cue was calculated as 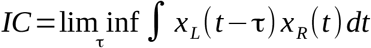 within the range of τ∈[-1,1] ms. The error based on IC was then calculated by comparing the target IC with the template IC:

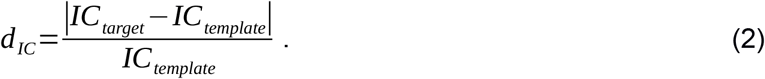

The errors for the other cues were calculated after filtering the target and template signals through a bank of fourth-order Gammatone filters with a regular spacing of one equivalent rectangular bandwidth [46]. The spectral excitation profiles were computed from the logarithm of the RMS energy within every frequency band [12, 47]. Audibility thresholds for band-limited signals were roughly approximated by assuming a sound pressure level of 70 dB and a within-band threshold of 20 dB. Further assuming stationary input signals, the spectral profiles were averaged over time, yielding spectral profiles as a function of frequency band, *p*(*f*). We chose this efficient approximation of auditory spectral profile evaluation as we observed in previous model investigations that a physiologically more accurate but less efficient approximation of the auditory periphery [48] lead to similar profiles and equivalent predictions of sound localization performance [37].

The interaural difference of the spectral profiles yielded IIDs as a function of frequency band, *IID*(*f*). For the ISSD cue, the model evaluated the standard deviation (SD) of *IID*(*f*) across frequencies and computed the negative difference of these deviations between the target and template, relative to the template deviation:

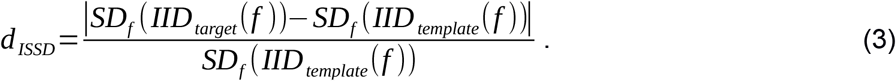

For the ISS cue, the absolute values of frequency-specific differences between the target and template IIDs were evaluated. Then differences smaller than 1 dB were set to zero and larger differences, Δ|*IID* (*f*)|, were normalized by the template IIDs and averaged across frequency bands, yielding:

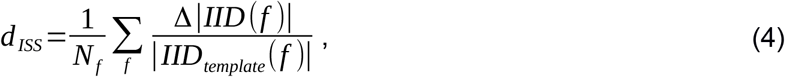

with *N*_*f*_ being the number of frequency bands.

The ITIT expectation error was calculated as the broadband deviation between target-to-template ratios of ITD and IID:

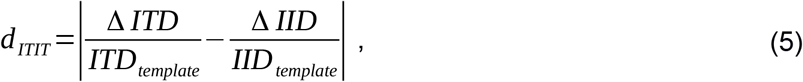

with Δ *ITD*=*ITD*_*target*_ − *ITD*_*template*_ and Δ *IID*=*IID*_*target*_ − *IID*_*template*_. Δ *ITD* and Δ *IID* smaller than ±20 µs and ±1 dB, respectively, were set to zero. The ITDs were derived from binaural signals following a procedure, in which the signals were low-pass filtered at 3 kHz and the ITD was the time lag that yielded maximum IC of the temporal energy envelope [49].

For the MSS and MSSD cues, positive spectral gradient profiles were derived exactly as in our previous work [37]. Briefly, first monaural spectral gradients were obtained by differentiating the excitation profiles (*p*(*f*)→ *p’*(*f*)) and softly restricting the value range by an elevated arctangent:

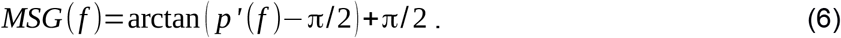

For the MSS cue, these gradients were then compared between the target and template separately for each ear by applying the same procedure as for the ISS metric, that is, calculating absolute target-to-template differences, normalizing differences larger than 1 dB (Δ|*MSG*(*f*)|) by the template gradients, and averaging those differences across frequencies:

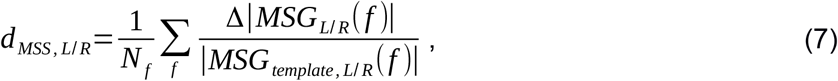

separately for the left and right ear as indexed by *L* and *R*, respectively. The MSS error metric was defined in analogy to ISSD:

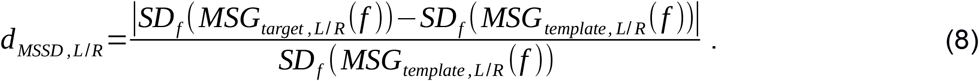

These unilateral error metrics were then combined according to a binaural weighting function [12, 13], effectively increasing the perceptual weight of the ipsilateral ear with increasing lateral eccentricity:

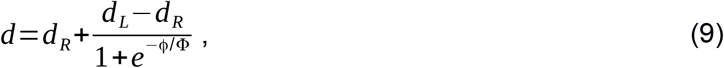

with ϕ∈[-90°, 90°] denoting the lateral angle (left is positive) and Φ=13°.

### Mapping to externalization ratings

A sigmoidal mapping function scaled by 2*e*_*range*_, shifted by *e*_*offset*_, and slope-controlled by a sensitivity parameter *S*_*cue*_ was used to map the metrics of expectation error *d*_*cue*_ to externalization ratings *e*_*cue*_ :

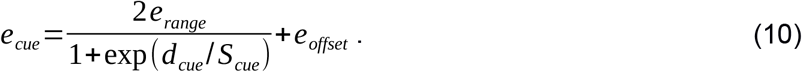

The numerator was doubled because the rating scale used in previous experiments was usually one-sided with respect to the reference sound, i.e., listeners were asked to rate only decreases and not increases of perceived externalization.

The mapping function in Eq. (10) contains one free model parameter, *S*_*cue*_, inversely related to the slope of the function mapping changes in the deviation metrics to changes in the externalization ratings. The mapping sensitivity is denoted by 1/ *S*_*cue*_ because larger 1/ *S*_*cue*_ yield steeper mapping functions that project smaller errors to smaller externalization ratings. This cue- and experiment-specific sensitivity was obtained by minimizing the squared simulation error, which was defined as the mean squared differences between actual and simulated externalization ratings (normalized scales). For the minimization, we applied the Nelder-Mead simplex (direct search) method (fminsearch, Matlab Optimization Toolbox, The Mathworks Inc.).

### Decision stage

For the WSM, optimal weights for scaling the contribution of a cue to the final rating were obtained by minimizing the simulation error. We used the same optimization technique as used for *S*_*cue*_. Weights smaller than 0.005 were considered as negligible and were set to zero.

For a fair comparison across our simulations of the decision strategies, the same number of model parameters was considered in the optimization. For the dynamic strategies, the mapping parameters were optimized to best fit the data across all experiments. For the weighted-sum strategy, the mapping parameters were fixed and corresponded to those from single-cue simulations, and the individual summation weights were optimized to best fit the data across all experiments.

### Considered studies

We simulated results of five previous headphone experiments [17, 33–35]. The pool of experiments was a compromise of data availability and the degree to which the test conditions isolate specific cues.

In Exp. I and II, Hartmann and Wittenberg [33] synthesized the vowel /a/ with a tone complex consisting of 38 harmonics of a fundamental frequency of 125 Hz, yielding a sound limited up to 4750 Hz. This sound lasted for 1 s and was presented via headphones and filtered with listener-specific HRTFs corresponding to 37° azimuth (0° elevation). The original phase responses of the HRTFs were maintained (the same applies also to the other studies). The HRTFs were manipulated at the frequencies of individual harmonics and the listeners rated the perceived externalization of those sounds presented via headphones on a labelled four-point scale (ranging from “0. The source is in my head” to “3. The source is externalized, compact, and located in the right direction and at the right distance”). The loudspeaker used to measure each listener’s HRTFs remained visible throughout the experiment. In Exp. I, the tone magnitudes up to a certain harmonic of a complex tone were set to the interaural average, effectively removing spectral IIDs up to that harmonic’s frequency. In Exp. II, instead of removing the original spectral IIDs, they were maintained. The *ipsilateral* magnitude spectrum was flattened up to a certain harmonic and these changes were compensated by shifting the contralateral magnitudes. This procedure maintained the original spectral IIDs but modified the monaural spectral profiles. For modeling the average results of these experiments, we used HRTFs from 21 exemplary listeners contained in the ARI database.

In Exp. III, Hassager et al. [17] investigated the effect of spectral smoothing on the auditory externalization of Gaussian white noises, band-limited from 50 Hz to 6000 Hz and lasting about 4 s. These sounds were filtered with listener-specific BRIRs (reverberation time of ∼0.2 s) in order to simulate sound sources positioned at azimuths of 0° and 50°. As independent experimental variable, Gammatone filters with various equivalent rectangular band-widths (ERBs) were used to spectrally smoothen either the direct path portion (until 3.8 ms) or the reverberation of the BRIRs. Filters with larger ERBs more strongly smoothed the shape of the magnitude spectrum and reduced the degree of externalization only when applied to the direct path. Smoothing applied to the reverberation affected the listeners’ externalization ratings only very little. Similar to Exp. I and II, listeners responded on a five-point scale where they perceived the sound as coming from (inside the head denoted as 1 and the visible loudspeaker position denoted as 5). Because the original BRIRs were not accessible, our model simulations were again based on the same 21 (anechoic) HRTFs from the ARI database and only addressed the strong effect of spectrally smoothing the direct path.

In Exp. IV, spectral smoothing was applied to Gaussian white noise bursts band-limited to higher frequencies from 1 kHz to 16 kHz [34]. Within that frequency range, listener-specific HRTFs used for binaural rendering were applied as measured (C = 1), as being spectrally flat (C = 0), or as something in between with reduced spectral contrast (C = 0.5). Pairs of about 0.5-s-long sounds were presented and the listeners were asked to judge whether the second sound was perceived closer or farther than the first sound. We evaluated only data from their “discontinuous trial” condition (of their Exp. II) with an inter-stimulus interval of 100 ms, which did not elicit an auditory looming bias and thus allowed to estimate absolute externalization ratings from these paired binary judgments by calculating mean relative frequencies of “farther” judgments. Two out of twelve listeners that unexpectedly perceived the spectrally flat sound (C = 0) as being farther than the individualized reference sound (C = 1) were not included in the present model analysis.

In Exp. V, Boyd et al. [35] used listener-specific BRIRs to simulate a talker positioned at 30° azimuth. The speech samples consisted of short sentences lasting about 3 seconds and provided sufficient speech energy up to 15 kHz. The study compared externalization ratings for in-the-ear (ITE) vs. behind-the-ear (BTE) microphone casings as well as broadband (BB) vs. 6.5-kHz-low-pass (LP) filtered stimuli at various mixing ratios with stereophonic recordings providing only an ITD but no IID. The listeners were asked to rate the degree of externalization against a preceding reference sound (1 s inter-stimulus interval) on a continuous scale with five labels (ranging from “0. Center of head” to “100. At the loudspeaker”). The reverberation time was 0.35 s. For our model simulations, original BRIRs were only available for three out of seven (normal-hearing) listeners.

Together the five experiments provide a good overview of the effects of different signal modifications on perceptual externalization. However, these studies were conducted in different laboratories and were applying slightly different methodologies, as summarized in Table 2. They differ, for instance, in whether they tested in an anechoic or reverberant environment and whether they provided visual information about the reference source or not. Such visual information may have contributed to a stabilized auditory externalization, but it does not seem to have degraded the effectivity of the signal modifications within the tested conditions. Factors like visual information and stimulus duration were ignored in our investigations (italic font in Table 2).

**Tab. 2.** Differences between considered studies with respect to methodological details and data availability. Visible refers to the visibility of a loudspeaker at the intended source location. Model simulations were either compared with the reported average results retrieved from the original publications (Avg.) or on the basis of individual results from N listeners. Modeled parameters are formatted in upright font, ignored parameters in italic font. For instance, the duration (Dur.) of stimuli was not considered in the model. In fact, non-spatial properties of the source signal besides its frequency bandwidth were only considered for Exp. I&II. In all other cases, the model was applied directly on the HRTFs or BRIRs.

| Exp. | Study | Signal | Bandwidth | *Dur.* | Azimuth | Reverb | Visible | Data |
| --- | --- | --- | --- | --- | --- | --- | --- | --- |
| I&II | [33] | Tone complex | 0.1 – 5 kHz | 1 s | 37° | none | yes | Avg. |
| III | [17] | *White noise* | 0.05 - 6 kHz | 4 s | 0°, 50° | *200 ms* | yes | Avg. |
| IV | [34] | *White noise* | 1 - 16 kHz | 0.5 s | 0°, ±90° | none | *no* | N = 10 |
| V | [35] | *Speech* | varied | 3 s | 30° | 350 ms | yes | N = 3 |

Despite Exp. IV, all the experiments used similar rating scales with the lowest rating representing the perception of a sound image located inside the head and the largest rating representing the perception of an image well-outside the head. In order to merge the responses from all studies into a single pool, we normalized the response scale of every individual experiment to externalization rating scores ranging from zero to one. Our normalized scores can be interpreted as the degree of externalization, mediating the egocentric distance percept [7], but not serving to determine the level of perceptual ambiguity per se. To hedge against the assumption of the rating scales being comparable across experiments, we also performed analyses with the externalization ratings treated as ordinal data. Those evaluations based on rank correlations yielded qualitatively similar but less distinguishable results. Hence, the here presented results focus on the normalized externalization scores.

## Results

### Individual cues: large differences across experimental conditions

We first investigated the ability of each individual cue to explain the effects of signal manipulations on auditory externalization as tested in the five experiments [17, 33–35]. Figure 2 shows the simulated externalization ratings together with the actual ratings from the original experiments. The frequency ranges of tested stimuli were classified as high or low with respect to the lower frequency bound around 5 kHz at which spectral cues can be induced by normal pinna morphologies.

**Fig. 2.**
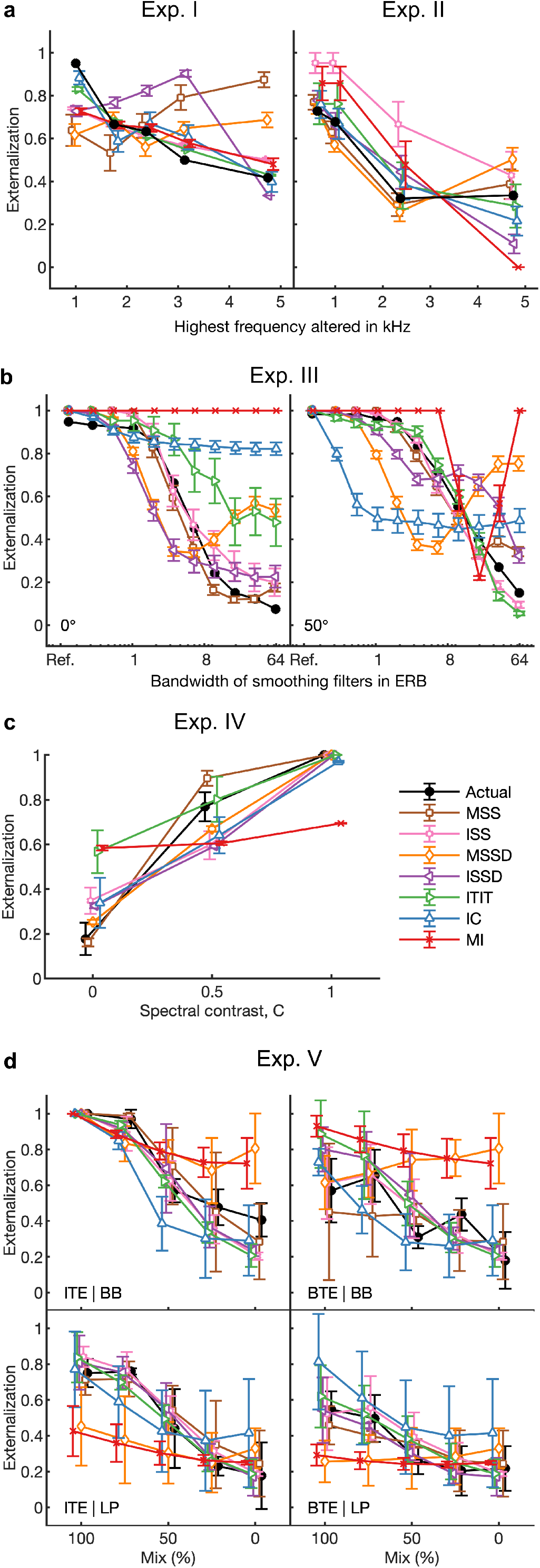
Externalization ratings: actual data from psychoacoustic experiments (closed circles) and simulations of the single-cue models (open symbols). **a**, Effects of low-frequency modifications tested by Hartmann and Wittenberg [33]. Exp. I: IID set to zero (bilateral average magnitude); actual data from their Fig. 7, average from N = 2. Exp. II: ipsilateral magnitudes flattened (IID compensated by contralateral magnitudes); actual data from their Fig. 8, average from N = 4. Simulated results for various cues, average from N = 21. **b**, Exp. III: effect of spectral smoothing of low-frequency sounds presented from various azimuths (left: 0°; right: 50°); actual data represents direct-sound condition from Hassager et al. [17], average from N = 7. Simulated N = 21. **c**, Exp. IV: effect of spectral smoothing in high frequencies as a function of spectral contrast (C=1: natural listener-specific spectral profile; C=0: flat spectra); actual data calculated from the paired-comparison data from Baumgartner et al. [34], N = 10 (actual and simulated). **d**, Exp. V: effects of stimulus bandwidth and microphone casing for various mixes between simulations based on listener-specific BRIRs (100%) and time-delay stereophony (0%); actual data from Boyd et al. [35], N = 3 (actual and simulated). ITE: in-the-ear casing; BTE: behind-the-ear casing; BB: broadband stimulus; LP: low-pass filtered at 6.5 kHz; Error bars denote standard errors of the mean.

#### Exp. I: Effect of IID modifications at low frequencies

Figure 2a (Exp. I) shows the effect of removing IIDs up to that harmonic’s frequency on the actual externalization ratings [33]. When simulating this experiment, the monaural spectral cues (MSS and MSSD) showed a non-monotonic relationship across the modified frequency ranges. Those simulated ratings largely diverged from the actual ratings especially for the largest frequency ranges (Fig. 2a, Exp. I) and resulted in large simulation errors (Fig. 3b, Exp. I). Further inspection showed that the modification of interaural averaging only marginally affected those monaural cues because at low frequencies the complex acoustic filtering of the pinnae is negligible and thus monaural spectral shapes are quite similar at both ears. In contrast, the broadband monaural cue (MI) was able to explain the actual ratings surprisingly well because the IID removal caused a small but systematic increase in sound intensity, being in line with the systematic decrease in externalization ratings.

**Fig. 3.**
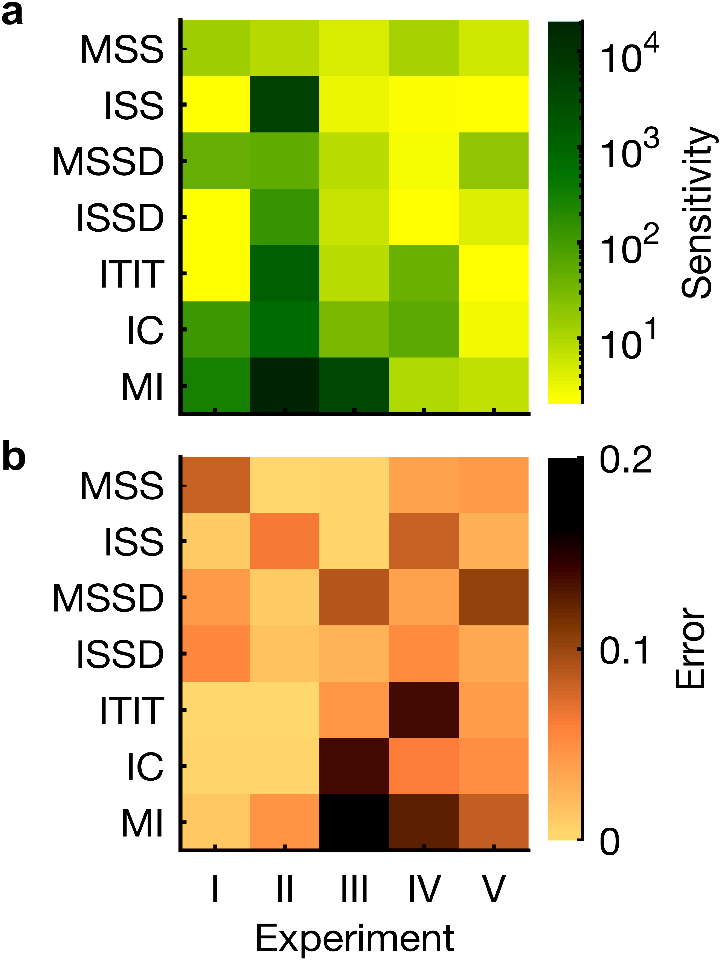
Optimization and performance of single-cue models. **a**, Cue-specific sensitivities used as optimization parameters. Higher values denote steeper mapping functions. **b**, Simulation errors as the RMS difference between the actual and simulated externalization ratings. Per experiment, the smallest error indicates the most informative cue.

Most interaural cues (ISS, ITIT, and IC) were able to explain the actual ratings very well. Differences in one interaural cue (IC) were, however, very small and thus required a very steep mapping function in order to become informative (indicated by a high sensitivity value in Fig. 3a, Exp. I). Simulations based on interaural spectral contrasts (ISSD) failed presumably because the evaluation of the standard deviation metric became unstable for spectral IID distributions strongly deviating from a normal distribution. Overall, the broadband interaural cue (ITIT) performed best with the minimum RMS simulation error of 0.06.

#### Exp. II: Effect of monaural modifications at low frequencies

In Exp. II, the interaural differences were maintained at the price of modifying the monaural spectral profiles. This resulted in a gradual distortion of perceived externalization (Fig. 2a, Exp. II).

In contrast to Exp. I, the simulations based on monaural spectral cues (MSS and MSSD) reflected the decrease in actual ratings very well, as indicated by RMS simulation errors for MSS as low as 0.05 (Fig. 3b, Exp. II). As expected, simulations based on spectral interaural cues failed because the degradation induced by flattening the ipsilateral spectrum was designed to maintain the original interaural features. Interestingly, the broadband interaural cues (ITIT and IC) were able to explain the effect of spectral manipulation on the ratings. As it seems, despite the compensation at the contralateral ear, the spectral flattening at the ipsilateral ear slightly modified both the IC and the discrepancy between the IID and ITD. While these small deviations from the template were able to explain the results from Exp. II, much larger sensitivities were required as compared to the simulations based on MSS and MSSD.

#### Exp. III: Effect of spectral smoothing at low frequencies

The spectral smoothing applied in Exp. III affected the auditory externalization of noise more gradually for the lateral as compared to the frontal direction (Fig. 2b) [17]. Again, only some of the cues were able to explain the systematic trend of externalization degradation. The stimulus manipulation affected some cues only marginally (IC, and MI), in a non-systematic manner (MSSD), or only for one of the two source directions (ITIT) whereas both monaural and interaural spectral shape cues (MSS and ISS) yielded more accurate simulations across conditions (RMS errors of 0.18 and 0.25, respectively, see Fig. 3b, Exp. III). Both cues yielded simulations consistent with the actual results in that they were insensitive to spectral smoothing below one ERB. It appears noteworthy that the monaural (MSS) outperformed the interaural (ISS) cue in this particular experiment, which has been used to promote an earlier modeling approach only based on that interaural cue [17]. Moreover, neither cue was able to explain why actual ratings were slightly below the maximum for the frontal reference sound (and bandwidths ≤ 1 ERB). Additional factors not represented in the model seem to be necessary to explain this lateral dependence of reference externalization.

#### Exp. IV: Effect of spectral smoothing at high frequencies

Focusing on the high-frequency range from 1 kHz to 16 kHz, where the pinnae induce the most significant directional spectral variations, Exp. IV concordantly showed that spectral smoothing degrades externalization perception (Fig. 2c) [34]. Most accurate simulations were obtained with monaural spectral cues (MSS and MSSD). The results for the other cues were mixed. For example, the two broadband cues ITIT and MI were hardly affected by the spectral smoothing and were not able to explain the actual results very well. While ITIT provided good results for C=0.5, it failed for signals lacking any spectral cues (C=0), predicting unreasonably high externalization ratings of over 0.5. The MI cue failed in all conditions. On the other hand, IC, the other broadband cue, showed relatively high simulation accuracies (Fig. 3b, Exp. IV). So there was no clear pattern for broadband cues. The simulations based on interaural cues were also ambivalent: ISS resulted in large simulation errors, while IC and ISSD showed smaller errors, indicating that some interaural information might have contributed.

Combining the results from Exp. III and IV shows that the effects of spectral smoothing can be best explained through template comparison based on monaural spectral shapes (MSS) and that this is independent of both the particular smoothing method and frequency range.

#### Exp. V: Bandwidth limitation and sensor displacement

Exp. V presented broadband speech samples from a mid-left position and compared externalization ratings for different sensor positions, stimulus bandwidths, and mixing ratios for a gradual removal of IIDs. The simulated ratings showed much variability across cues and conditions (Fig. 2d). The broadband BTE condition caused the most distinct spectral modifications and was particularly informative about the explanatory power of considered cues. Most cues were able to follow the actual ratings quite well (except MSSD and MI). On average across conditions, the simulation results suggest that expectations based on interaural spectral templates (ISS, simulation error of 0.16, see Fig. 3b, Exp. V) have been most relevant to the listeners in this experiment.

#### Individual cue summary

The overall picture of simulation errors suggests that there is no single cue able to explain the externalization ratings for all experiments but that there is a small set of cues on which listeners seem to base their externalization ratings. These cues correspond to the evaluation of monaural and interaural spectral features (MSS and ISS, respectively) as well as the broadband interaural disparities between ITD and IID (ITIT). This is consistent with previous observations showing that both interaural and monaural changes affect externalization perception.

The monaural cue (MSS) yielded best simulations in three out of five experiments as well as on average across the five experiments (average simulation error of 0.18). The evaluation of this cue focuses mainly on positive gradients in the spectral profile as motivated by physiological findings in the cat dorsal cochlear nucleus [50, 51]. To assure that our results were not confounded by a suboptimal choice of gradient sign, we repeated all simulations also with two different model configurations either favoring negative or considering both positive and negative gradients (technically, we implemented this by either switching the signs or removing the ±π/2 shifts in Eq. (6), respectively). We found that the average simulation error increased to 0.24 (for negative gradients only) and to 0.23 (for both negative and positive gradients), consolidating our choice of focusing the evaluation on positive spectral gradients.

The sensitivity parameters (shown in Fig. 3a) used to scale the mapping function from cue-specific error metrics to externalization scores were optimized separately for every cue and experiment. The separate optimization was reasoned by the limited quantitative comparability of subjective externalization ratings because of various factors such as differently trained subjects, different contexts, different experimental procedures, and other methodological differences. Nevertheless, the optimization procedure yielded somewhat similar sensitivity parameters for the same cue across experiments whenever that cue was informative as indicated by small RMS simulation errors.

In summary, we found considerable variance in cue-specific simulation accuracy. Simulations based on a single monaural cue (MSS) turned out to be most potent in explaining the set of the considered experiments. However, the best cue clearly depended on the experiment and simulations only based on that particular cue would fail in situations similar to Exp. I. This indicates that the auditory system does not simply rely on a single cue and leads to our main research question of which perceptual decision strategy is used to combine cues in different listening situations.

### Decision strategy: static weighting outperforms dynamic selection

We aimed to determine which decision strategy best explains externalization ratings across all experiments without an a-priori knowledge about the context. According to the principle of the static decision strategy (WSM), the simulated listener derives the final externalization rating from a linearly weighted combination of single-cue ratings (Fig. 1b). For that strategy, the weights were obtained from an optimization procedure minimizing the simulation errors. The dynamic decision strategies, that is, WTA, MTA, and LTA, selected the largest, median, and smallest simulated externalization rating, respectively, obtained across the considered cues.

By doing so, the cue weights become binary: the cue providing the largest, median, or smallest externalization rating is considered as the total rating, all other cues are finally ignored.

Overall, the static cue combination (WSM) outperformed the dynamic strategies, as indicated by significantly lower simulation errors (Fig. 4a) and higher rank correlations (around 0.9 for WSM, 0.8 for MTA and LTA, and 0.5 for WTA). It simulated the externalization ratings based on the weighted sum of mainly two spectral cues: a monaural cue (MSS) weighted by about 60% and an interaural cue (ISS) weighted by 40%, while the other cues contributed only marginally (Fig. 4b). The contribution of the interaural cue is in line with previous findings in the context of Exp. III, for which only interaural cues have previously been used to explain externalization ratings [17]. Across other experiments, as shown here, the monaural cue becomes essential.

**Fig. 4.**
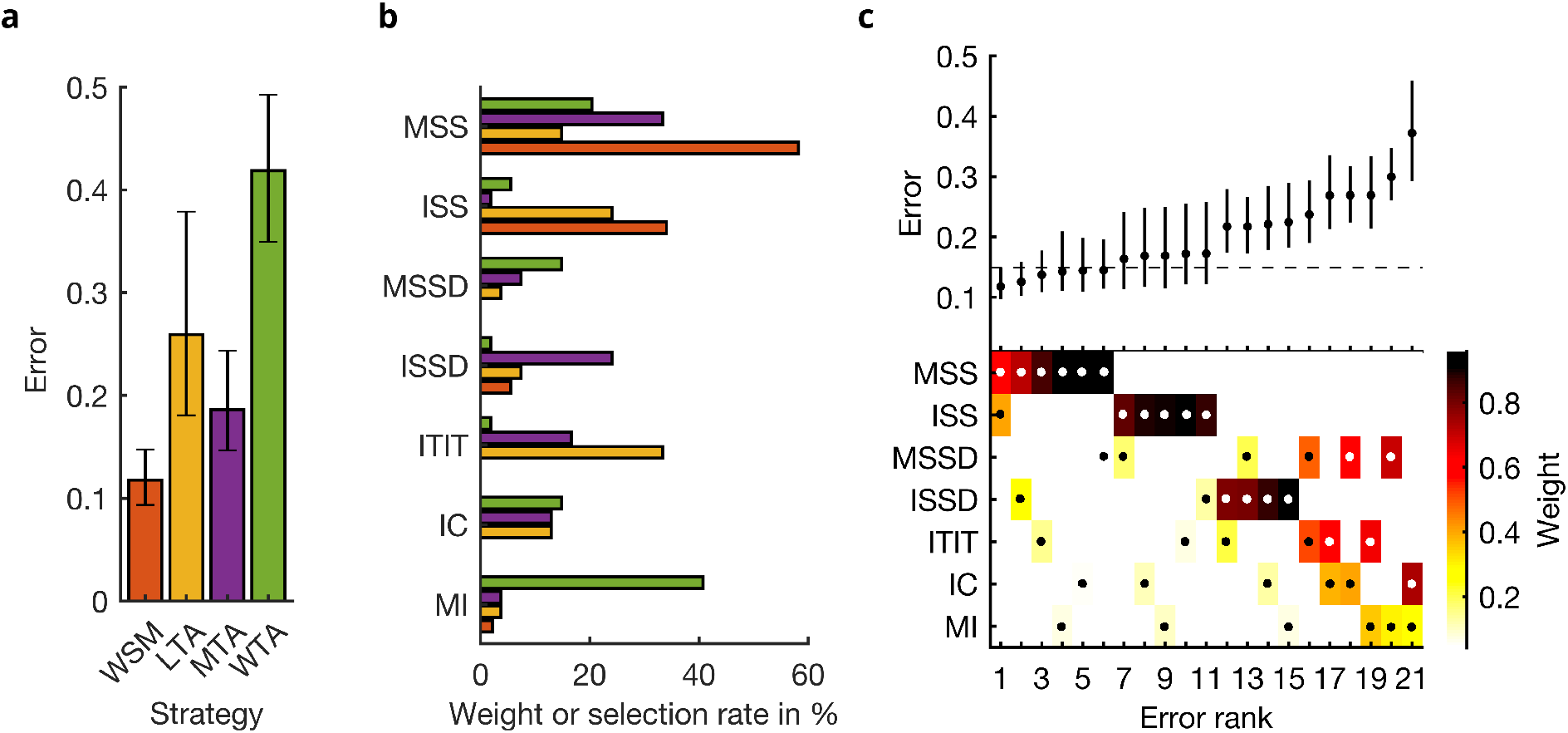
Simulation errors for different decision strategies and cue combinations show that static combination (WSM) based on monaural and interaural spectral shape cues (MSS, ISS) performs best. **a**, RMS simulation errors for different strategies and pooled experimental data, N = 54. Error bars denote 95% confidence intervals estimated via bootstrapping (1000 resamples). WSM: weighted-sum model; L/M/WTA: looser/median/winner takes all. **b**, Individual cue contributions. Cue abbreviations defined in Tab. 1. **c, Top:** Simulation errors for pairwise combinations of considered cues. Dashed line shows the border of significant difference to the best pair (MSS and ISS). **Bottom**: Considered cue pairs (highlighted by dots) with their respective weights (encoded by brightness).

Among the dynamic strategies, the conformist approach (MTA) performed better than all the other selection approaches (Fig. 4a). The perfectionist approach (LTA) showed intermediate performance and the minimalist approach (WTA) performed particularly poorly, effectively equivalent to a simulated chance performance of 0.42 RMS error. The cues selected by the dynamic strategies were more diverse than the weights of the static strategy (WSM, Fig. 4b). In the MTA strategy, the monaural cue (MSS), also favored by the static strategy, played an important role, suggesting that it provides the most consistent ratings across all considered cues. In the LTA strategy, broadband interaural cues (ITIT) were most frequently selected, which is in accordance with our single-cue simulation results where this particular cue also provided good results for some conditions. The large variance in simulation errors in particular for LTA (error bars in Fig. 4a) further indicates that deviations in single cues fail to explain externalization ratings in some of the experimental conditions even if the selected cue is allowed to change with condition.

### Cue relevance: both monaural and interaural spectral shapes matter

Our analysis suggests that under spatially static listening conditions, auditory externalization is mainly based on two cues. However, most of the cues co-vary to some degree, and thus the relevance of a particular cue as a contributor to the cue combination may be strongly affected by the joint consideration of dependent cues. To investigate the effect of such interdependency, we simulated listeners using the static decision strategy (WSM) with only two cues but all possible combinations.

The results, sorted by increasing simulation error, are shown in Fig. 4c. As expected there is a considerable variance in the simulation errors obtained by different pairs, with an order of magnitude between the best and worst pair. The condition considering the previously favored cues (MSS and ISS) yielded the smallest simulation error, confirming our findings from simulations considering more than two cues.

The results underline the importance of both cues by two means. First, all pairs including the monaural cue (MSS) were ranked highest and the simulation error remained small regardless of the other considered cue. As long as that cue was involved, the simulation errors did not increase significantly (remained within the confidence intervals of the best condition). Thus, the MSS cue seems to be the most important cue for auditory externalization under static listening conditions. Second, by not considering that monaural cue, the error increased (with the mean above the confidence intervals of the best condition), but was still not significantly different to the best condition, as long as the spectral interaural cue (ISS) was considered in the modeling. In fact, the simulation errors significantly increased (p < 0.05, compared to the best condition) as soon as neither the monaural (MSS) nor the interaural (ISS) cues of spectral shape were considered. Other spectral cues that only evaluate the amount of spectral contrast (MSSD and ISSD) instead of its shape failed to explain perceived externalization.

As a hypothetical negative benchmark that was not tested in any of the considered studies, we briefly investigated also a perfectly diotic noise condition without any HRTF filtering. For such a listening situation, the model following the WSM strategy under the assumption of a frontal incidence angle predicts an externalization score of around 0.2. It being larger than zero is mainly caused by the fixed contribution of ISS that detects only small deviations to the template because the ISS is generally very small for positions on the median plane.

## Discussion

In order to uncover the essential cues and the decision strategy underlying auditory externalization perception in various static listening situations, we developed a template-based modeling framework and applied it to a set of psychoacoustic experiments investigating how spectral distortions degrade the auditory externalization percept. Our results suggest that a static, weighted combination (WSM) rather than a dynamic selection (LTA, MTA, WTA) of monaural and interaural spectral cues (MSS and ISS, respectively) drives perceptual decisions on the degree of auditory externalization. Hence, although listeners are sensitive to many individual cues, in spatially static conditions, their externalization perception seems to be driven by a static combination of only few cues with fixed perceptual weights.

### Dominant cues

The two major cues favored by our model selection procedure are both based on spectral shapes. Monaural [52] as well as interaural [53] spectral cues are known to be informative about absolute distance in the acoustic near-field of a listener. Those two cues are also well-known to be important for and complementary in the process of sound localization.

Monaural spectral shapes (MSS) constitute localization cues within sagittal planes, i.e., including both the vertical and front/back dimension [44]. The current understanding of this cue is that it resembles the processing of monaural positive spectral gradient profiles [12, 37], in line with electrophysiological measurements in the dorsal cochlear nucleus [50]. Concordantly, reducing the contribution of negative spectral gradients here turned out to improve the simulations of perceived externalization.

Interaural intensity differences are very well established as a cue for auditory lateralization [15, 16]. Given the inherent frequency selectivity of the auditory periphery it is reasonable to assume that relative differences to an internal reference are evaluated on a frequency-specific level [17], as implemented in the ISS cue of our model. On the other hand, the accessibility or predictability of such an interaural spectral evaluation must be limited somehow in the auditory system because otherwise it would not make sense to consider the monaural counterpart at all for spatial inference, as this cue intrinsically suffers from ambiguity with the source spectrum [54]. Moreover, the ISS-induced overestimation of externalization ratings for a hypothetical diotic noise condition may indicate that the currently used definition and/or incorporation of the ISS cue is incomplete. For instance, it could be that ISS is contributing only at lateral directions where IIDs are significantly larger than zero.

### Template matching in view of predictive processing

In order to assess incoming cues most efficiently and rapidly, theories of predictive processing assume a hierarchical propagation of error signals representing differences between internal predictions (templates) and sensory evidence on increasingly complex matters, while errors are weighted by the expected sensory reliability for that matter [55]. In view of auditory externalization, the ultimate goal is to assign an allocentric spatial position to an auditory object and errors between expected and observed cues need to be weighted by the amount of information those cues offer for the inference of an externalized sound-source position.

It was unclear how flexibly those weights may be adjusted. Our results suggest that the weighting between spatial spectral cues, which are naturally persistent, remained fixed across the various experimental contexts. In contrast, monaural intensities, for instance, inform distance inference only on a relative scale with intrinsic ambiguity. The weighting of such naturally less persistent cues was outside the scope of this study but would be expected to highly depend on the listening context. It would be interesting to target the short-term adaptability of cue weights by varying their reliability in future experiments.

### Limitations and directions for future investigations

Reverberation is another less persistent but usually omnipresent cue. Reverberation smears the spectral profile and decorrelates the binaural signal, effectively increasing the variance of interaural cues and affecting the reliability of spectral cues. The degree of externalization increases with the amount of reverberation if the reverberation-related cues are consistent with the listener’s expectations about the room acoustics [56]. In parallel to the present work, we investigated also the impact of situational changes of the amount of reverberation on externalization perception by extending the here promoted framework to reverberation-related cues [36]. There, we introduced a weighting factor that accounts for the reduced reliability of spectral cues with increasing amount of reverberation-related cues while not affecting the relative weighting between the two spectral cues.

The externalization models proposed here and there do not explicitly consider interactions with perceived incidence angle [57–60]. However, the strong overlap of cues between the perceptual externalization and directional sound localization suggests a strong overlap of the underlying perceptual processes. In our simulations, template comparisons were only performed for the reference direction, ignoring that there might be a strong match to a template from another direction yielding strong externalization at that other perceived location. For example, spectral cue changes may have elicited changes in the perceived polar angle within a sagittal plane. On the other hand, in most of the considered studies, the experimental paradigms involved rating of externalization against a fixed reference stimulus and, thus, one can assume that listeners were able to ignore directional localization changes. Nevertheless, joint assessment of externalization and directional localization in future behavioral experiments is needed to further our understanding of the interdependency between those two perceptual processes.

As for directional localization, the perceptual inference underlying auditory externalization can also adapt to changes in the naturally very persistent spectral cues [61]. Given a fixed weighting of those cues in the model framework, this adaptivity should be solely grounded on the generation of new expectations, i.e., an updating of templates. How such new templates evolve and compete with existing templates would be an exciting objective for future research.

A more substantial extension of the model framework will be required in order to explain perception in dynamic listening situations. Especially self-generated movements like head rotations are known to elicit strong expectations on its acoustic consequences and drastically degrade externalization if not being fulfilled [57, 59, 62]. In contrast to reverberation-related cues, the spectral cues studied here are strongly affected by such movements. Internal generative models are considered to constantly shape the listener’s expectations by means of either explicit or implicit predictive codes [55]. Embedding the here proposed model in a larger predictive coding framework and using Bayesian inference to account for sensory uncertainty [63, 64] may pave the road for future work more deeply investigating the short-term dynamics and multisensory context in spatial hearing.

From a technical perspective, our present model successfully predicts auditory externalization ratings under spatially static conditions without prior knowledge about the listening context. This is a feature often requested by engineers working on audio rendering and human interfaces within the context of augmented and virtual reality [65–67], while considering the spatially static condition as a worst case scenario regarding sensitivity to spectral distortions. In particular, the model can be applied to efficiently assess perceived externalization in a variety of acoustic applications relying on spectral cue modifications such as passive binaural audio playback of static sources over headphones or hearing aids [7]. A future extension to spatially dynamic situations will further extend the range of potential applications. To this end, a concept of modeling sound localization in dynamic and active listening situations has recently been proposed [64].

To simplify the application and further development of the here proposed model, and to guarantee reproducibility of results, implementations of both the model (baumgartner2021) and the model simulations (exp_baumgartner2021) are publicly available as part of the auditory modeling toolbox (AMT, version 1.0, https://www.amtoolbox.org) [68].

## Conclusions

Perception of externalized events depends on the alignment of observed auditory cues with the internal expectations of the listener. In order to uncover the most essential cues and the strategy used by the listener to integrate them, we compared various model types in their capability to explain five experiments focussing on the effects of spectral distortions within a spatially static environment. Most accurate simulations were obtained for the combination of monaural and interaural spectral cues with a fixed relative weighting (about 60% monaural, 40% interaural). In contrast, the dynamic selection of cues depending on its situational magnitude in expectation error was less successful in explaining the experimental results. Together, this suggests that at least in spatially static environments, listeners jointly evaluate both monaural and interaural spectral cues to perceptually infer the spatial location of the auditory object. Further, the proposed model framework appears useful for predicting externalization perception in binaural rendering applications.

## Acknowledgements

We want to thank Bill Withmer for kindly providing parts of the original data from Boyd et al. (2012) as well as Henrik Gert Hassager, Barbara Shinn-Cunningham and H. Steven Colburn for fruitful discussions. This work was supported by the Austrian Science Fund (FWF, grant J 3803-N30 to R.B.), Oculus VR LLC, and the European Commission (grant 691229).

## Author Contributions

R.B. conceptualized and implemented the research, analyzed the results, and wrote the original draft; R.B. and P.M. interpreted the results and revised the manuscript.

